## Supplementary Figure s6 for "Mechanical force of uterine occupation enables large vesicle extrusion from proteostressed maternal neurons"

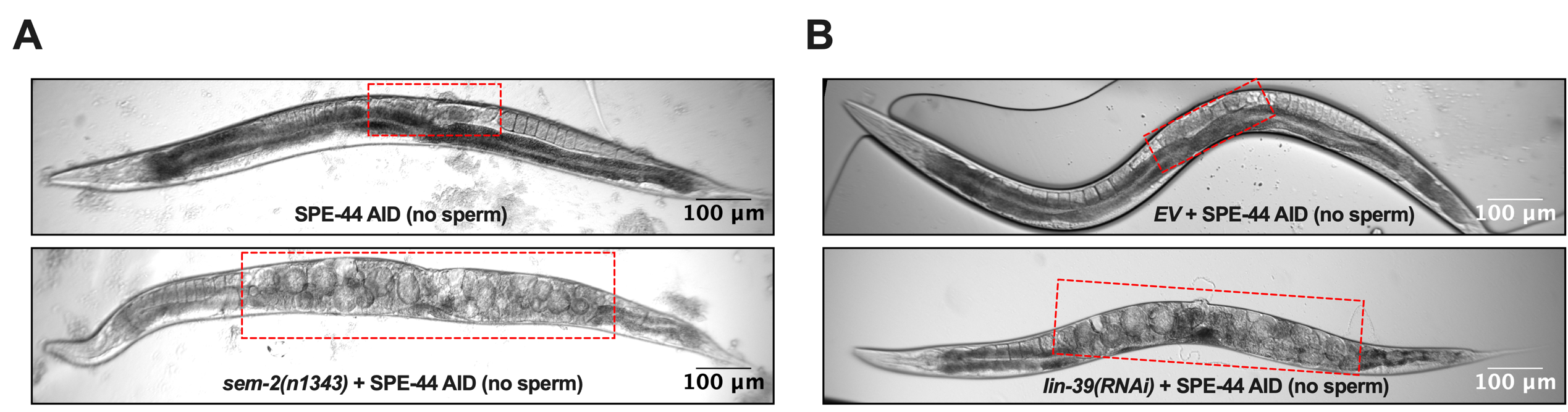

**Figure s6. Representative pictures of oocytes retention (red rectangle) in the uterus of Adult day 2 hermaphrodite.**

A. Strain ZB4749 (top): fxIs1[P*_pie-1_*::TIR1::mRuby] zdIs5[P*_mec-4_*::GFP] I; bzIs166[P*_mec-4_*::mCherry] II; *spe-44(fx110[spe-44::degron]*) IV vs. ZB4953 (bottom): *sem-2(n1343)* fxIs1[P*_pie-1_*::TIR1::mRuby] I; bzIs166[P*_mec-4_*::mCherry] II; *spe-44(fx110[spe-44::degron]*) IV. 1 mM auxin treatment induces no sperm status to both strains.
B. Strain ZB4749: fxIs1[P*_pie-1_*::TIR1::mRuby] zdIs5[P*_mec-4_*::GFP] I; bzIs166[P*_mec-4_*::mCherry] II; *spe-44(fx110[spe-44::degron]*) IV treated with either control empty vector (EV) (top) or *lin-39(RNAi)* (bottom). 1 mM auxin treatment induces no sperm status.
